# Quick and effective approximation of *in silico* saturation mutagenesis experiments with first-order Taylor expansion

**DOI:** 10.1101/2023.11.10.566588

**Authors:** Alexander Sasse, Maria Chikina, Sara Mostafavi

## Abstract

To understand the decision process of genomic sequence-to-function models, various explainable AI algorithms have been proposed. These methods determine the importance of each nucleotide in a given input sequence to the model’s predictions, and enable discovery of *cis* regulatory motif grammar for gene regulation. The most commonly applied method is *in silico* saturation mutagenesis (ISM) because its per-nucleotide importance scores can be intuitively understood as the computational counterpart to *in vivo* saturation mutagenesis experiments. While ISM is highly interpretable, it is computationally challenging to perform, because it requires computing three forward passes for every nucleotide in the given input sequence; these computations add up when analyzing a large number of sequences, and become prohibitive as the length of the input sequences and size of the model grows. Here, we show how to use the first-order Taylor approximation to compute ISM, which reduces its computation cost to a single forward pass for an input sequence. We use our theoretical derivation to connect ISM with the gradient of the model and show how this approximation is related to a recently suggested correction of the model’s gradients for genomic sequence analysis. We show that the Taylor ISM (TISM) approximation is robust across different model ablations, random initializations, training parameters, and data set sizes.

## Introduction

Deep learning models have become the preferred tool to analyze the relationship between genomic sequence and genome-wide experimental measurements such as chromatin accessibility ^1,2^, gene expression^3–5^, 3D chromatin conformation ^6–8^, and other molecular data modalities ^9–11^. To understand the models’ decision processes, and extract the learnt genomic features, various explainable AI algorithms have been developed ^12–14^. These methods estimate the importance of each nucleotide in an input sequence to the model’s predictions.

The most commonly used algorithm to interpret genomic sequence-to-function models is in *silico* saturation mutagenesis (ISM)^15^. ISM is very straightforward to implement and biologically highly interpretable. It can be intuitively compared to performing *in vivo* saturation mutagenesis experiments ^16^, as ISM computes the change in the model’s prediction as a function of a change in a single nucleotide. More formally, given a trained sequence-to-function model, at every position *l* along an input sequence of length *L* the reference base *b*_*0*_ is replaced by one of the other three alternative bases *b*_*v*_∈{A,C,G,T | *b*_*v*_ ≠ *b*_*0*_} one at a time, and the model’s predictions on the alternative sequences recorded. Thus, to compute the ISM profile for a sequence of length L, 3xL forward passes are required. The differences between the predictions of these variant sequences and the prediction from the “reference” (initial) sequence is then used to define the impact or importance of the reference base and each alternative base along the sequence.

It is hoped that when applied to increasingly accurate sequence-based deep learning models, ISM can aid in solving for a comprehensive *cis* regulatory grammar and in some cases replace laborious and expensive *in vivo* saturation mutagenesis experiments ^4,9,17^. However, as the state-of-the-art models continue to model larger input sequences (e.g., >100Kb), it is becoming computationally prohibitive to apply ISM. Here, we study the effectiveness of a first-order Taylor approximation ^18^ to compute ISM values along massive sets of sequences using the model’s gradient from a single forward pass with each sequence. We show that TISM approximations speed up computations of ISM values by a factor L times 3 divided by the batchsize. TISM derived attribution maps highly resemble attribution maps from ISM, more than the models’ gradient as it is. We also derive that TISM represents the theoretical link between a recently proposed correction of the model’s gradient for investigation of genomic sequences and attribution maps derived from ISM. Importantly, we show that TISM values are robust across different models, random initializations, training parameters, and data set sizes.

## Background

To approximate the value of a complex function *f* at input *s*, Taylor’s approximation linearly decomposes the function value *f(s)* at *s* into the value of the function at nearby position *s*_*0*_, given by *f(s*_*0*_), and the derivative of the function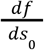at *s*_*0*_ multiplied by the difference between the position of interest *s* and *s*_*0*_ .

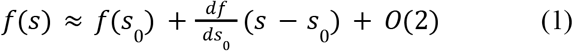

Where O(2) represents the second order term that is truncated in the linear approximation. In the case of sequence-based deep learning model producing the function *f*, the gradient 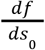 can also be represented as the finite difference from the reference *s*_0_ to a sequence with a single nucleotide substitution from *b*_0_ to *b*_1_ at position *l*, denoted by *s* (*l, b*_0_→*b*_1_ ).

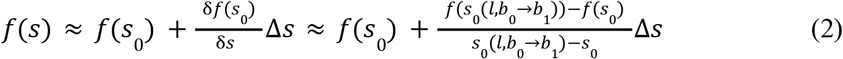

In the case where the finite distance δs to approximate the gradient is equal to the distance from the reference sequence Δs, the numerator and denominator cancel each other out and we are left with the ISM value given by Equation (3).

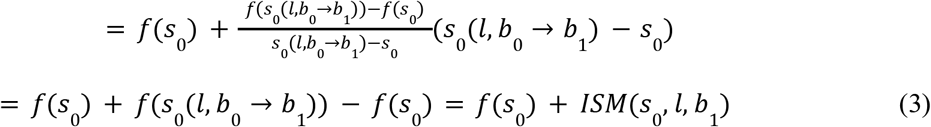

Thus, Equation (3) shows that ISM can be understood as the linear effect from an approximation of the deep learning model *f* with respect to a surrogate variable Δ*s* = *s* − *s*_0_ . In practice, ISM values are used in two ways: (1) as a per-nucleotide value that indicates how much the prediction changes if the reference base is replaced by the specific variant; (2) as attribution maps which indicate how important each nucleotide is for a model’s prediction. To generate attribution maps from ISM values, practitioners subtract the mean of at each position *l* from its ISM values to get attributions per base-pair *A*_*ISM*_*(s*_*0*_, *l, b*_*v*_).

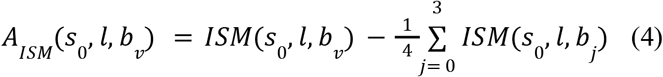

Regulatory motifs are usually identified from the values of these attribution maps at the reference base. They represent how important the present nucleotide is for the model’s predictions akin to common measures of per-nucleotide sequence conservation.

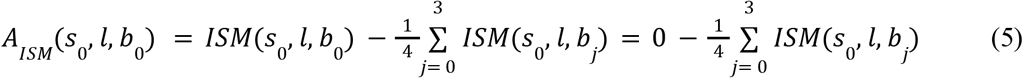

While ISM is very easy to implement, it is computationally costly, and so users often resort to using “gradient-times-input” to indicate how important the given nucleotide is for a model’s prediction ^4,9^. Gradient-times-input is considerably less computationally taxing, because it uses a single pass through the model to simultaneously approximate the importance of every nucleotide in the input sequence. Specifically, during model training, the gradient with respect to the parameters is computed automatically in every forward pass to enable parameter updates with backpropagation. Therefore, the gradient with respect to the input is available for “free” from just a single forward pass through the network. For model interpretation, gradient-times-input simply uses the gradient at the reference base, indicating whether it is beneficial for the model to either “change”, or keep the base at this position.

Here, we propose to instead use the gradient to approximate ISM using a first-order Taylor approximation. By equating *f(s)* in equations (1) and (3), we see that ISM can be approximated from the model’s gradients:

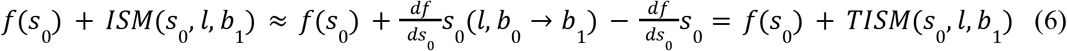

where TISM denotes the first-order Taylor approximation to ISM. Applying this to a one-hot encoded input in which the reference base *b*_*0*_ a position *l* is replaced (set *b*_*0*_ from 1 to 0) by an alternative base *b*_*1*_ (set b_1_ from 0 to 1), we can see that the ISM at *l,b*_*1*_ is equal to the gradient with respect to the reference sequence *s*_*0*_ at base *l,b*_*1*_ minus the gradient at base *l,b*_*0*_.

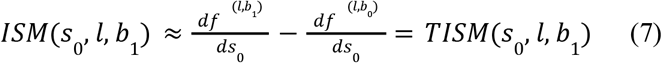

Using this relationship allows us to quickly approximate the per nucleotide ISM values from the gradient of the input sequence using only a single forward pass through the model. This is especially useful when applying ISM to long sequences, or for comparing regulatory motifs across many sequences. Distal regulatory elements are common in genomics, and estimating the correct effect size is key to determine their impact on gene regulation.

While gradient times input estimates the importance of the reference base from the gradient at the reference base 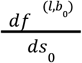, attribution maps derived from TISM correctly add the effect from the alternative bases to the attribution of the reference at position *l*.

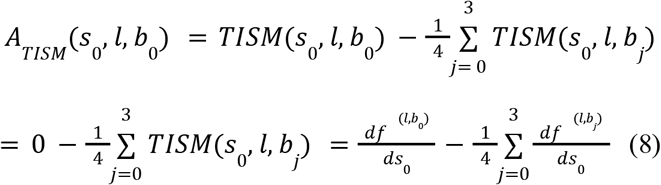

We note, recently Majdandzic et al. proposed the same correction for gradient-based attribution maps as TISM but motivated it from a geometrical point of view ^19^. Briefly, the authors suggested minimizing the impact of of-simplex gradient noise by removing the random orthogonal gradient component from the input gradient. Here, we showed that this correction simply represents an approximation of attribution maps that are derived from ISM. In addition, we show how these values can be biologically interpreted and how the gradients of the model are directly related to ISM values.

## Methods

To determine how well TISM reproduces the results computed by ISM under different conditions, we trained various versions of our well established AI-TAC model ^20^ on ATAC-seq counts within 286,000 open chromatin regions (OCRs, regions that were determined to be open in at least one cell type) across 81 mouse immune cells^21^. All models use a 251bp one-hot encoded sequence around the center of the ATAC-seq peaks and predict log_2_(x+2) transformed Tn5 cuts within the peak region. We train on OCR sequences from 16 out of 19 chromosomes, and use OCRs on chromosome 8 and 11 for validation, and those on chromosome 19 as an independent test set. We use early stopping and select our final model based on the highest mean Pearson’s correlation coefficient between model’s prediction of accessibility and the ground truth across cell types for each OCR sequence (computed on the validation set).

All the models use 298 kernels of length 19bp, with a GELU activation function, and softmax weighted mean pooling of size 2. The models apply 4 residual convolutional blocks with batch normalization, kernel size of 7, 298 kernels, and subsequent softmax weighted mean pooling of size 2. The resulting representation is flattened and condensed to a size 512 tensor with a linear layer, followed by two fully connected layers with GELU activation, before the model heads predict the log transformed counts for the 81 cell types in a multi-task fashion. All models use a dropout of 0.1 in all the fully connected layers. We use a mixture of MSE and mean correlation of OCRs across 81 cell types as a loss function and update the models’ parameters with SDG with 0.9 times momentum and a learning rate of 1e-5. The learning rate is exponentially warmed up in 7 epochs, and fine tuning is performed for five iterations on the best performing parameters with gradually reduced learning rates. All models use the forward and the reverse strand of the sequence and perform kernel specific max-pooling along the aligned forward and backward activations from both sequences, only forwarding the highest activation of a kernel from the two strands at a given position. In addition to the sequence centered at the ATAC-peak, we also perform data augmentation by including shifted versions of each sequence ^9^, where we randomly shift the genomic location of a given sequence by a number between -10 to 10 base pairs.

AI-TAC is a standard CNN model trained with correlation loss function. Because attribution methods we use here are agnostic to the specific model architectures, the results presented should generalize more broadly to other models and training datasets. However, we also repeated these experiments with various modifications to the model’s architecture to examine the generalizability of our results. Specifically, to test how concordant TISM values are to ISM values across different modeling choices/architectures, we trained different models and used ablation to investigate the effect of various modeling choices. First, we trained four versions of the above “baseline” model with different random initializations. Second, we trained the model on five different percentages of the training set to assess how training set set size influences TISM’s concordance to ISM. Third, we stopped training after 1, 2, 3, 5, 7, 11, 20, 60, 100, and 400 epochs and assessed how the concordance changes during training. Lastly, we performed ablation analysis of the baseline model as follows: 1) We trained solely using the MSE loss, 2) we used ReLU activation throughout the model, 3) we used an exponential activation function after the first convolution ^22^, 4) we used max-pooling instead of the weighted mean pooling, 5) we trained without the reverse complement sequence, 6) we trained without residual connections, 7) we trained without dropout, 8) we trained without batchnorm in the residual convolutional layers, 9) we used L1-regularization of the 298 kernels in the first layer, 10) we trained with AdamW instead of SGD, 11) we trained without randomly shifted sequences, and 12) we trained a shallow CNN that only uses one convolutional layer, 70 bp wide weighted mean pooling, which is flattened and then directly given to the linear prediction head.

To empirically determine the speedup of TISM over ISM, we trained three additional model architectures that used sequences of length 1,000bp, 5,000bp, and 20,000bp as input. For these models, we used the same number of convolutional layers but adjusted the size of the softmax weighted mean poolings to account for the larger sequence windows. We then determined the time to compute ISM and TISM values for models with four different input lengths and for one model for four different numbers of sequences. Specifically, to compute ISM values, we measured the total time to generate a set of one-hot encoded variant sequence tensors that contain a single base-pair change to the original sequence, make predictions with the model using a batch size of 20, and finally generate the ISM tensor from these predictions. For TISMs, the measured time includes the forward pass through the model, backpropagation through the model to the input sequence to obtain the gradient, and finally subtraction of the gradient at the reference base from each position to get TISM values. All the measurements were performed on a single NVIDIA RTX A4000 GPU with 16GB Memory.

## Results

We used the trained modified version of our previously published model AI-TAC (see Methods) and evaluated the concordance between ISM and TISM on the per nucleotide effects in a given input sequence. The model takes a DNA sequence of 251bp around the ATAC peak (OCR) as input and predicts the normalized accessibility (i.e. the number of Tn5 cuts within 250 bp around the ATAC peak, corrected for sequencing depth) of that peak across 81 different cell types in a multitask fashion. The model was trained on 286,000 OCRs and ISM and TISM values were computed for 9,158 OCR sequences for all 81 cell types (i.e 741,798 attribution maps each). In all evaluations below, we solely use regions from chromosome 19 which were entirely left out during model training and validation.

In our baseline model, we observed an average correlation value of 0.7 between TISM and ISM values (**Figure 1a**). Encouragingly, 87% of TISM profiles computed from test regions had a correlation value of at least 0.6. Visual inspection confirmed the high concordance between ISM and TISM profiles across different cell types and suggests that both methods detect the same motifs and predict similar changes to their effect across cell types (**Figure 1b**). Next, we confirmed our theoretical derivation and compared TISM to the gradient as a popular alternative to ISM. TISM’s correlations to ISM are consistently higher than those of the gradient itself (**Figure 1c**). We also compared concordance between the mean effect per base from TISM and ISM versus the concordance of the mean effect from ISM to gradient-times-input (**Figure E1**). While the correlation of the mean effect per base from TISM to ISM is also consistently higher, gradient times input’s correlation to ISM is closer than the correlation of gradients across all four bases.

**Figure 1.**
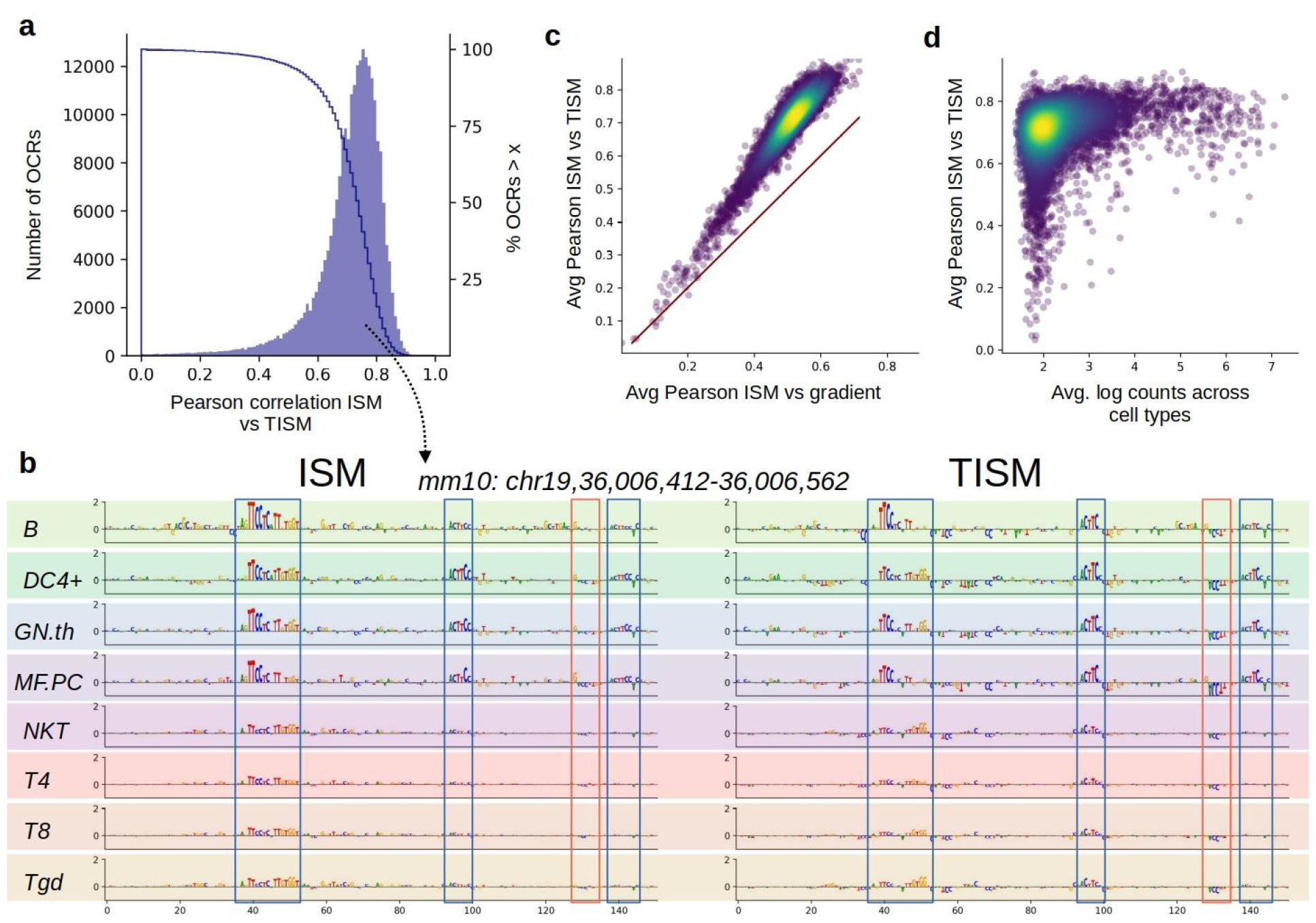
Comparison between ISM and Taylor approximated ISM. **a)** Histogram and cumulative percentage of Pearson’s correlation between ISM and TISM of 9158 chromatin regions across 81 cell types in the test set (Mean = 0.7, Median =0.73). **b)** Attributions of reference base from ISM and TISM for ‘Atac.peak_251276’ at chr19:36,006,362-36,006,612 across 8 cell types with differential chromatin accessibility. Selected peak’s ISM and TISM correlate 0.75 across all 81 cell types. Blue squares indicate consistent motifs between TISM and ISM. Red squares indicate motifs that are inconsistent (here only present in TISM) **c)** Average Pearson correlation of OCRs across all cell types between ISM and the Gradient (x-axis) versus the average correlation between ISM and Taylor corrected gradient (TISM). **d)** Correlation between peaks’ mean log accessibility and the correlation between ISM and TISM is R=0.29.

We examined the sequences with lower correlation between TISM and ISM and noted that for most part, these correspond to regions of low average predicted chromatin accessibility and high coefficients of variations of predicted counts (**Figure 1d, E2a**). We did not observe a relationship between model performance and ISM to TISM correlation (**Figure E2b**). When we measured the running times for both methods, we confirmed the theoretical speedup of TISM over ISM (**Table 1a, b**). When we measured the speedup for different numbers of sequences, TISM was on average ∼160 times faster than ISM (**Table 1a**). This is consistent with theoretical values from using a batch size of 20 to compute the ISM values (3 times 1,000bp divided by batch size of 20). TISM exerts its real value for long sequences, where its speedup improves from 25 times to 8,000 times for sequences of length 251bp to sequences with length 20,000bp (**Table 1b**). We note that this speedup is 2.5 times larger than expected with a batch size of 20. This is likely due to other choices in our implementation of ISM, such as generation of alternative sequences, that take up extra time (see Methods).

**Table 1.**
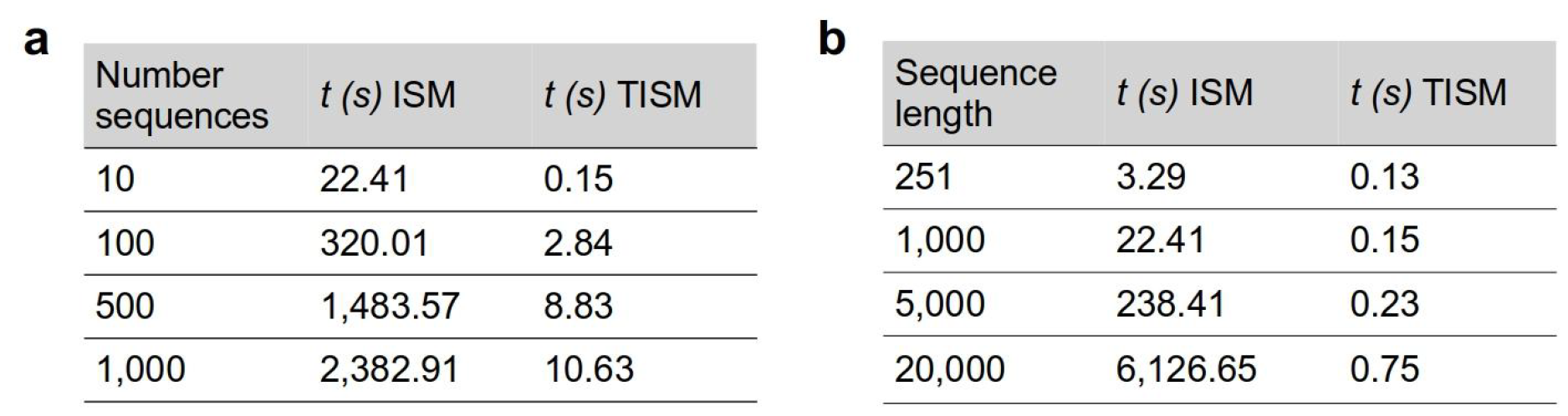
Comparisons between run times of ISM and TISM. For all models and experiments, a batch size of 20 was used to perform forward passes through the models **a)** Run times to generate ISM and TISM values for 10, 100, 500, and 1,000 sequences of length 1,000bp. **b)** Run times of models that take as input sequences of length 251, 1,000, 5,000, and 20,000bp to compute n=10 ISM and TISM values.

| **a** |  |  | **b** |  |  |
| --- | --- | --- | --- | --- | --- |
| Number sequences | $t$ (s) ISM | $t$ (s) TISM | Sequence length | $t$ (s) ISM | $t$ (s) TISM |
| 10 | 22.41 | 0.15 | 251 | 3.29 | 0.13 |
| 100 | 320.01 | 2.84 | 1,000 | 22.41 | 0.15 |
| 500 | 1,483.57 | 8.83 | 5,000 | 238.41 | 0.23 |
| 1,000 | 2,382.91 | 10.63 | 20,000 | 6,126.65 | 0.75 |

To determine how robust these approximations are across models, we trained our baseline model from four random parameter initializations and computed the Pearson correlation between TISM and ISM profiles from all four models (**Figure 2a**). On average, the correlation of ISM profiles from two separate model runs is 0.58. TISM profiles behave similarly, with an average correlation of 0.52: However, TISM and ISM profiles correlate on average with Pearson R=0.7 when they are from the same model, showing that TISM is more concordant with ISM than ISMs between different model trainings.

**Figure 2.**
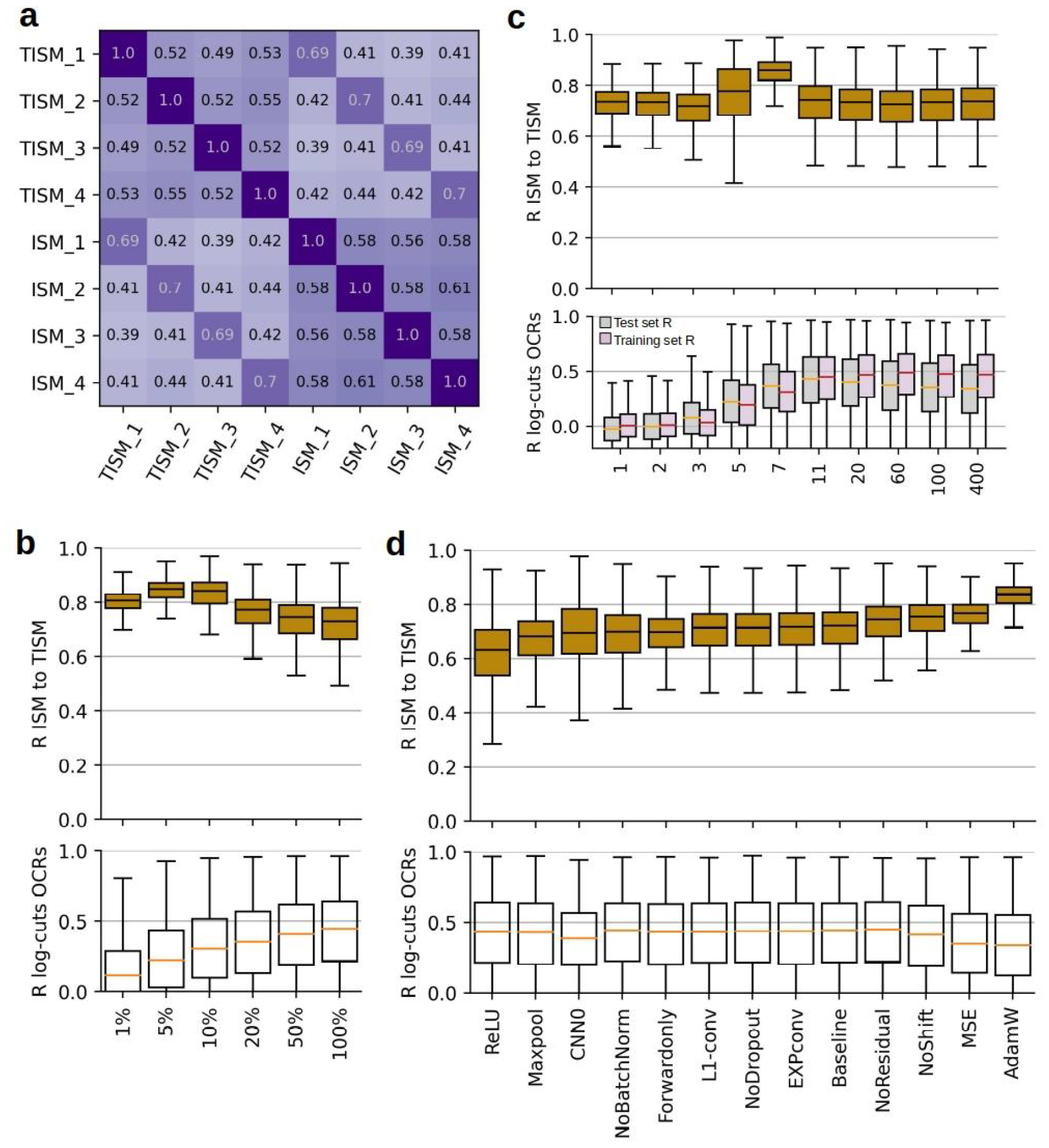
Comparison between ISM and Taylor approximated ISM across models. **a)** Average correlation between ISM and TISM values across 3,000 regions and 20 cell types from four random model initializations. The AI-TAC model was trained four times from different random parameter initialization. **b)** TISM to ISM correlation for 3,000 test set regions across 81 cell types for six models trained on different percentages of the training set. Bottom shows the Pearson correlation between predicted and measured log-counts of OCRs in the test set across 81 cell types **c)** TISM to ISM correlation for different ablations of the baseline model. Pearson correlation between predicted and measured log-counts of the twelve models for 9158 chromatin regions across 81 cell types.

Next, we examined the concordance of TISM profiles with decreasing size of training data. To do so, we sub-sampled OCRs to create a couple of different smaller training sets (1%, 5%, 10%, 20% and 50%). For each training set size, we trained a model on the randomly selected subset of data points and evaluated the correlation between TISM and ISM on the same 3,000 peaks across 81 cell types (243,000 ISM profiles; **Figure 2b**). Surprisingly, we observe that the correlation between ISM and TISM is higher for models that were trained on a subset of the data points, with the largest concordance between the two at 5% of the data points, or 14,300 training data points. Simultaneously, the predictive performance of the model is decreasing as expected (**Figure 2b bottom**). From visual inspection, we observe that models trained on smaller data sets are missing regulatory motifs that are present when trained on larger data sets (**Figure E3a**, blue frames). Additionally, the TISMs from smaller data sets are also missing negative motifs that are not concordant with the ISM effects (**Figure 1b, E3a**, red frames). We hypothesize that these discordant motifs in models trained on larger data sets are the result of non-linear effects that TISMs cannot account for by the first-order Taylor approximation, even within proximity of only single nucleotide change.

To investigate this further, we looked at the evolution of concordance between TISM and ISM during model training. We trained a model for 400 epochs and assessed the concordance between ISM and TISM after 1, 2, 3, 5, 7, 11, 20, 60, 100, and 400 epochs (**Figure 2c**). As expected, the test set performance (mean Pearson correlation R of OCRs across cell types) is increasing until 11 epochs and then slightly decreases afterwards while the training performance continues to increase. We note that the model is not overfitting as strongly as we would expect normally. We assume that this is due to our architectural choices, the Pearson correlation loss across cell types and the sequence shifting in particular. At the beginning of model training, we observe that the concordance between ISM and TISM profiles is similar to our fully trained baseline model while its predictions are still random (mean_Test_ = -0.02, mean_Train_ = 0.01). Interestingly, the concordance between ISM and TISM increases after three epochs and reaches its optimum with a mean Pearson of 0.85 at epoch seven, when model performance is slightly less than optimal (mean_Test_ = 0.37). Once the model reaches its optimum performance in the test set at epoch 11 (mean_Test_ = 0.41) the concordance has decreased back to an average 0.71. Additional overfitting (epoch 400: mean_Test_ = 0.34, mean_Train_ = 0.45) does not affect the concordance between TISM and ISM (mean concordance 0.71). We hypothesize that the increase in correlation before reaching the optimal model performance is the result of the model learning linear relationships which the first-order approximation can well represent. However, afterwards the model potentially starts learning non-linear effects that increase its performance but reduce the concordance between ISM and TISM.

Lastly, we trained twelve different models, each with a single ablation to the baseline model. We used these models to study the impact of model architecture on the accuracy of TISMs approximations (**Figure 2d**, Methods). Most training and architectural choices did not affect the concordance between ISM and TISM (i.e. exponential activation, no dropout, L1 on kernel, forward strand only). While removing batchnorm, and max-pooling did not affect the performance of the model, both choices slightly decreased the concordance between TISM and ISM (mean_NoBatchnorm_=0.68, mean_Maxpool_=0.66). The shallow CNN *CNN0* performed slightly worse in performance but has the same mean concordance as the baseline model (Mean = 0.7).

We observed the worst concordance between ISM and TISM from a model that used ReLU activations across the network (mean_ReLU_=0.61), while this choice did not result in worse predictions. On the other hand, using AdamW (mean=0.82), MSE loss (mean=0,76), and no sequence shifting (mean=0.74) during model training, resulted in worse performance but higher concordance between ISM and TISM. Visual inspection of ISM and TISM profiles generated by a model trained with AdamW suggest that the updates with weight decay lead to smaller motif effects (**Figure 1b, E3b**, blue frames) and less varying gradients outside the well-defined motifs (**Figure 1b, E3b**, red frames). Since non-linear effects are rare in the data, weight decay, in addition to reducing the size of the linear effects, likely removes rare non-linear effects entirely, leading to more concordant TISM but less accurate predictions.

## Discussion

Here, we provide the missing theoretical link between ISM and gradient-based interpretation methods for sequence-to-function models, which we call Taylor approximated ISM (TISM). We use TISM to generate 741,798 sequence attribution maps of length 251bp for 22 models, and assess the concordance between the computationally effective approximation and the directly computed values across sequences. We find that TISM is highly concordant with ISM values and possesses higher correlation to ISM than ISMs between different model initializations. Concordance between the two gets worse for model architectures that use ReLU and max-pooling layers which potentially make it more challenging to accurately compute the model’s gradients. Counterintuitively, we also observe that models trained on fewer data points, model architectures with worse predictive performance, or not fully trained models possess higher concordance between ISM and TISM. We hypothesize that this can be explained by these underperforming models having learned only simple motif grammar that misses interaction terms between bases. Further exploiting this observation we hypothesize that one could use the discordance between TISM and ISM to detect sequences that harbour strong base-pair interactions.

Issues with understanding attribution maps result from limited understanding of what these values mean in a functional sense. ISM is biologically interpretable but can become computationally challenging for large sets of long sequences that are processed by deep networks. While other backpropagation based methods can help with this, their values are often harder to interpret and therefore hard to compare across, positions, sequences, and models. The recently developed geometrical correction to the model’s gradient by Majdandzic et al. ^19^ shows empirical and anecdotal evidence for improving motif identification but does not provide a theoretical link to other attribution maps or functional explanation of what these geometrically corrected motif values represent.

Here, we show how one can very simply approximate ISM from the model’s gradient. Approximating ISM enables the analysis of both large sets of sequences and long sequences. TISM’s strength especially comes through for long sequences (e.g., > 20kb), and therefore it is extremely useful to detect, extract, and compare regulatory motifs across sequences and tasks ^4^. While not as accurate as FastISM ^23^ or Yuzu ^24^ (because these are not approximations), TISM, in contrast, is applicable to any network written in any code base, any number of sequences, and only requires a few lines of code to turn the model’s gradient into TISM values.

We show that the majority of TISM (89%, >0.58) values correlates well above ISM values from different model initializations, suggesting that TISM is sufficient to understand the model’s learned regulatory grammar and predict effects of sequence variants across different loci.

## Extended data figures

**Figure E1.**
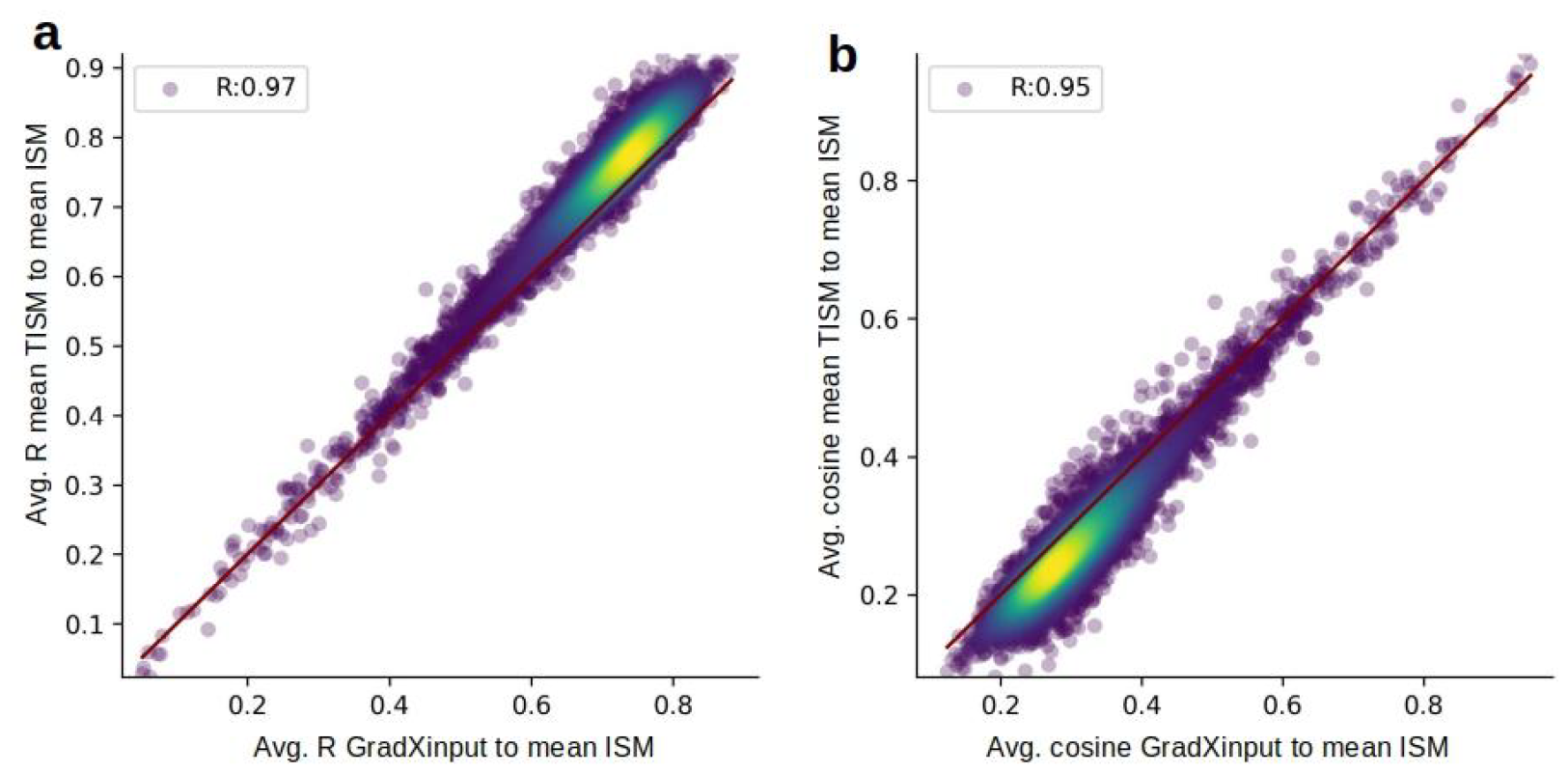
Comparison between mean TISM and gradient times input. **a)** Average Pearson correlation of OCRs across all cell types between mean effect of TISM to mean effect of ISM (y-axis) compared to average Pearson from gradient times input (x-axis) to the reference attribution of ISM. **b)** Average cosine distance correlation of OCRs across all cell types between reference attribution of TISM reference attribution of ISM (y-axis) compared to average cosine distance from gradient times input (x-axis) to the mean effect of ISM.

**Figure E2.**
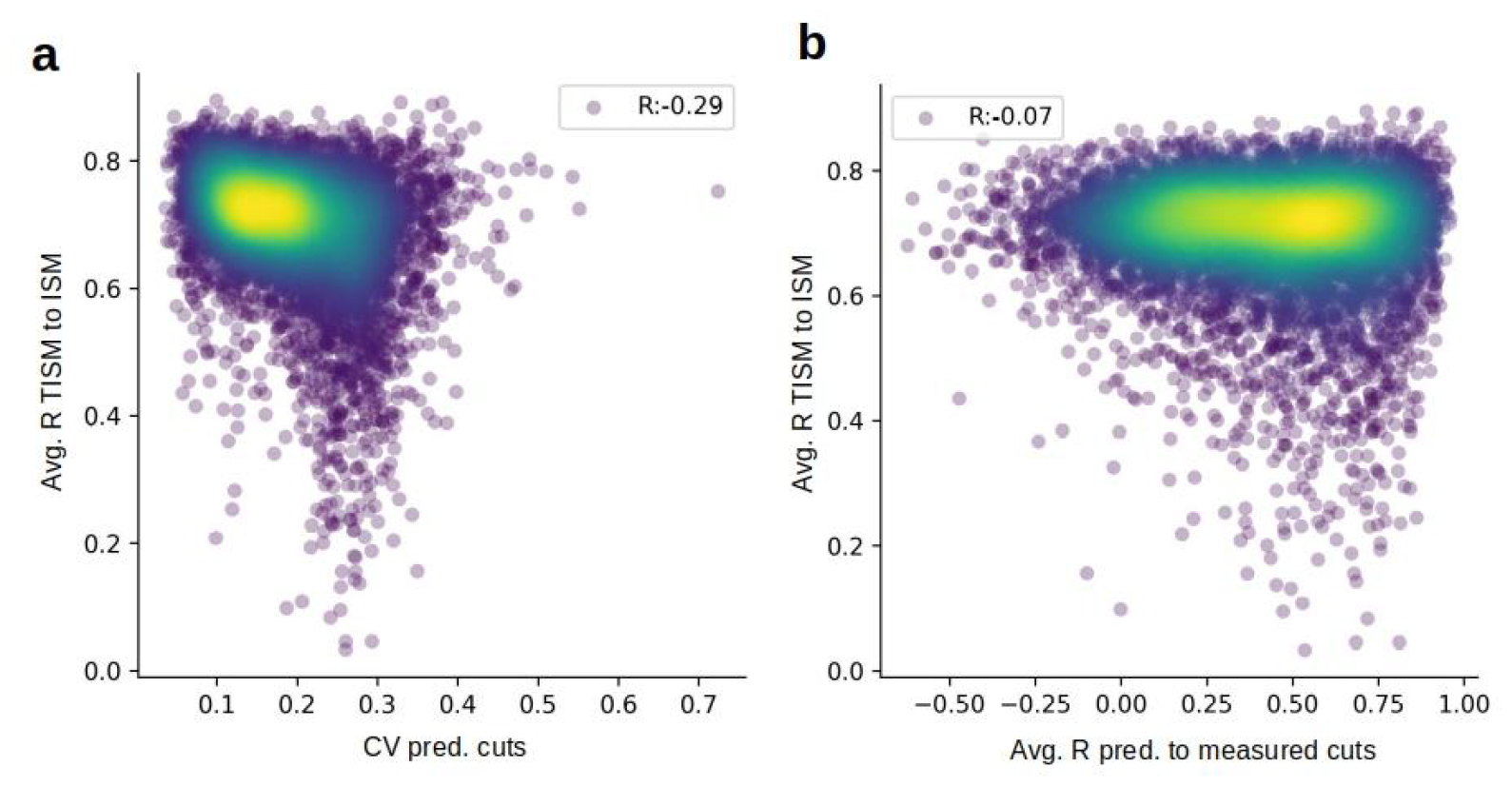
Relationship between concordance of TISM and ISM and model predictions. **a)** Relationship between peaks’ coefficient of variation of predicted log cuts and the correlation between ISM and TISM. **b)** Relationship between model performance of OCRs (Pearson correlation R across cell types) and the correlation between ISM and TISM.

**Figure E3.**
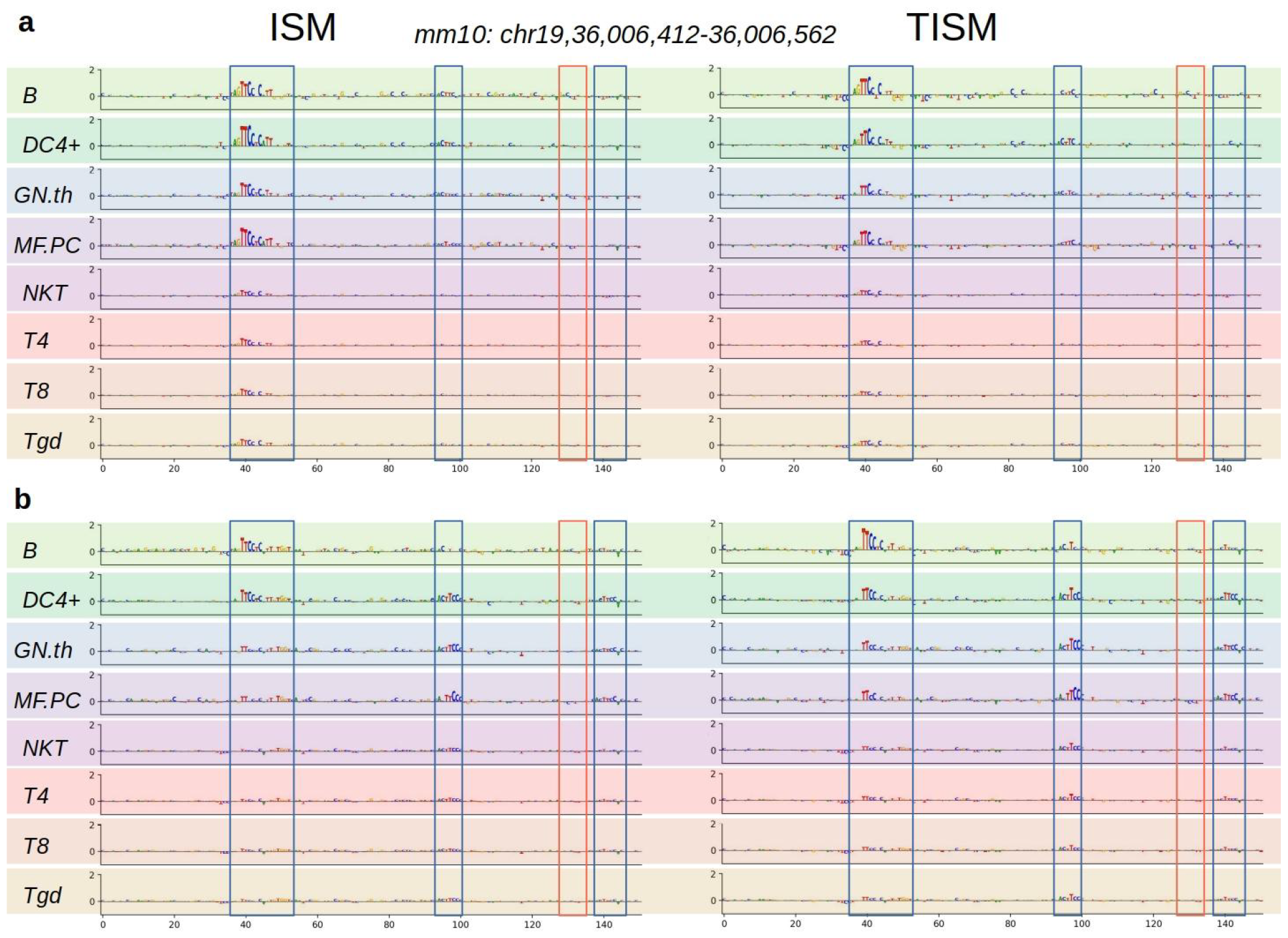
Comparison between reference attributions from ISM and TISM at chr19:36,006,362-36,006,612 across 8 cell types with differential chromatin accessibility. Blue squares indicate consistent motifs between TISM and ISM. Red squares indicate motifs that are inconsistent (here only present in TISM) **a)** ISM and TISM from the AI-TAC model that was trained on 5% of the data (see Figure 2b, Methods). **a)** ISM and TISM from the AI-TAC model that was trained with AdamW (see Figure 2d).

